# Finding human gene-disease associations using a Network Enhanced Similarity Search (NESS) of multi-species heterogeneous functional genomics data

**DOI:** 10.1101/2020.03.11.987552

**Authors:** Timothy Reynolds, Jason A. Bubier, Michael A. Langston, Elissa J. Chesler, Erich J. Baker

## Abstract

Disease diagnosis and treatment is challenging in part due to the misalignment of diagnostic categories with the underlying biology of disease. The evaluation of large-scale genomic experimental datasets is a compelling approach to refining the classification of biological concepts, such as disease. Well-established approaches, some of which rely on information theory or network analysis, quantitatively assess relationships among biological entities using gene annotations, structured vocabularies, and curated data sources. However, the gene annotations used in these evaluations are often sparse, potentially biased due to uneven study and representation in the literature, and constrained to the single species from which they were derived. In order to overcome these deficiencies inherent in the structure and sparsity of these annotated datasets, we developed a novel Network Enhanced Similarity Search (NESS) tool which takes advantage of multi-species networks of heterogeneous data to bridge sparsely populated datasets.

NESS employs a random walk with restart algorithm across harmonized multi-species data, effectively compensating for sparsely populated and noisy genomic studies. We further demonstrate that it is highly resistant to spurious or sparse datasets and generates significantly better recapitulation of ground truth biological pathways than other similarity metrics alone. Furthermore, since NESS has been deployed as an embedded tool in the GeneWeaver environment, it can rapidly take advantage of curated multi-species networks to provide informative assertions of relatedness of any pair of biological entities or concepts, e.g., gene-gene, gene-disease, or phenotype-disease associations. NESS ultimately enables multi-species analysis applications to leverage model organism data to overcome the challenge of data sparsity in the study of human disease.

**Availability and Implementation:** Implementation available at https://geneweaver.org/ness. Source code freely available at https://github.com/treynr/ness.

**Author summary:** Finding consensus among large-scale genomic datasets is an ongoing challenge in the biomedical sciences. Harmonizing and analyzing such data is important because it allows researchers to mitigate the idiosyncrasies of experimental systems, alleviate study biases, and augment sparse datasets. Additionally, it allows researchers to utilize animal model studies and cross-species experiments to better understand biological function in health and disease. Here we provide a tool for integrating and analyzing heterogeneous functional genomics data using a graph-based model. We show how this type of analysis can be used to identify similar relationships among biological entities such as genes, processes, and disease through shared genomic associations. Our results indicate this approach is effective at reducing biases caused by sparse and noisy datasets. We show how this type of analysis can be used to aid the classification gene function and prioritization of genes involved in substance use disorders. In addition, our analysis reveals genes and biological pathways with shared association to multiple, co-occurring substance use disorders.

## Introduction

A fundamental problem in computational biology is the identification and characterization of the biomolecular basis of complex disease. However, human genetic and genomic studies are necessarily sparse due to the costs and the complexity of studying the human population at a molecular level [1]. In contrast, model organism studies are quite information rich due to the availability of specimens in controlled experiments and the tremendous array of technologies available for acquisition of genomic data [2]. Supplementation of human datasets with model organism experiments represents a powerful means with which to study complex disease. However, cross-species data integration presents a major technical challenge, both at the level of identifying similarity among disease related features, and assessing the conservation of biomolecular mechanisms, particularly gene regulatory relations.

Historically, characterization and comparison of biological concepts, such as disease and gene products, has relied heavily on descriptive observations (e.g., symptomatology). Modern approaches use controlled vocabularies and ontologies to document observable phenotypic characteristics associated with the underlying disease [3, 4]. This approach, coupled with semantic similarity methods for example, can be used to compare diseases [5]. One difficulty in using solely curated or semantic data sources, is the tendency for curatorial processes to produce annotations that are biased toward well-studied species and processes, resulting in an uneven distribution of annotations [6, 7]. Furthermore, gene annotations are affected by the idiosyncrasies of experimental systems and assays [8] which may impact downstream analysis.

More recently, methodologies integrating genomic datasets and interaction networks have been proposed, leading to refined comparison of biological entities based on large-scale, experimentally derived data. Typically, these approaches utilize protein-protein interactions (PPI) [9] and functional networks [10] as a supplement for sparse gene annotation datasets. Network-based integration has been used to uncover disease-disease relationships [11], identify gene-disease modules [12], and re-prioritize putative, pathogenic genetic variants [13]. Among integrative network methods, random walks and network propagation have been particularly useful for applications such as gene-disease prioritization [14], inferring gene-phenotype relationships [15], and disease classification [16]. For complex disorders such as autism [17] and cancer [18], these network-based approaches have been successful at uncovering and prioritizing the underlying genetic components.

Overall, integrative network analysis has shown great potential for identifying the genetic mechanisms of disease. However, most approaches utilize datasets from a single species, primarily humans, rather than exploiting the vast amount of extensive model organism experiments that characterize the depth and breadth of intrinsic biological mechanisms of disease processes. Additionally, most approaches lean on highly curated but sparse data sets representing only a specific data type, e.g., PPIs, co-expression networks, curated gene-disease annotations, leaving a wealth of experimentally derived, heterogeneous functional genomics data unused or under-utilized.

In the present study, we describe a methodological approach, Network Enhanced Similarity Search (NESS), for improving the accuracy and sensitivity of similarity comparisons among biological entities that leverages these additional data types. Briefly, we aggregate and harmonize a number of heterogeneous graph types including ontologies and their annotations, biological networks, and bipartite representations of experimental study results across species. We employ diffusion metrics, specifically a random walk with restart (RWR), to estimate the relations among entities in the graph and to make data-driven comparisons. The advantage of this methodology is three-fold: first, it enables the integration of heterogeneous data types, genomic data, and big data stores to alleviate biases inherent in unevenly studied concepts and the idiosyncrasies of experimental systems; second, human experimental results can be supplemented with model organism studies by accounting for orthologous gene relations across many species; and finally, a graph representation of genomic feature relations allow complex biological mechanisms to be modeled, including gene regulatory mechanisms (e.g., expression quantitative loci, epigenetic marks, and 3-D genomics). We show this approach is resilient to sparse and noisy datasets, and outperforms other concept comparison methodologies when using expert-curated resources such as the Gene Ontology (GO) and Kyoto Encyclopedia of Genes and Genomes (KEGG) as a ground truth. We assess the performance of this method at prioritizing genes involved in complex disease–a subset of substance use disorders–and compare these results with the state of the art. Additionally, NESS is coupled with a graph permutation framework that can be used to empirically assess the statistical significance of any results.

We make NESS freely available as a software package which can be retrieved from https://github.com/treynr/ness. We also provide an implementation of this tool for use within the GeneWeaver resource (https://geneweaver.org) which allows NESS to use a background of over 100,000 heterogeneous functional genomics datasets.

## Materials and methods

### Data integration and heterogeneous network assembly

In this work we provide a means to assess the similarity of biological concepts through their genetic underpinnings. This is possible through cross-species data integration of heterogeneous functional genomics data. First, graph-based structures are retrieved and constructed from experimental, functional genomics and public resource data sets. A cross-species heterogeneous graph is generated from these sources. (Fig 1). Currently the network supports 15 different nodes (Table 1) and is amenable to new data types. Nodes are connected to one another using edge relationships such as ontology annotations, gene set contents, and genetic interactions. Edges are typically undirected but in some cases, such as when embedding ontology graphs and their annotations within the network, directed edges are used. The network is also amenable to incorporation of edge weights; for simplicity, only unweighted edges are used in this analysis.

**Figure 1.**
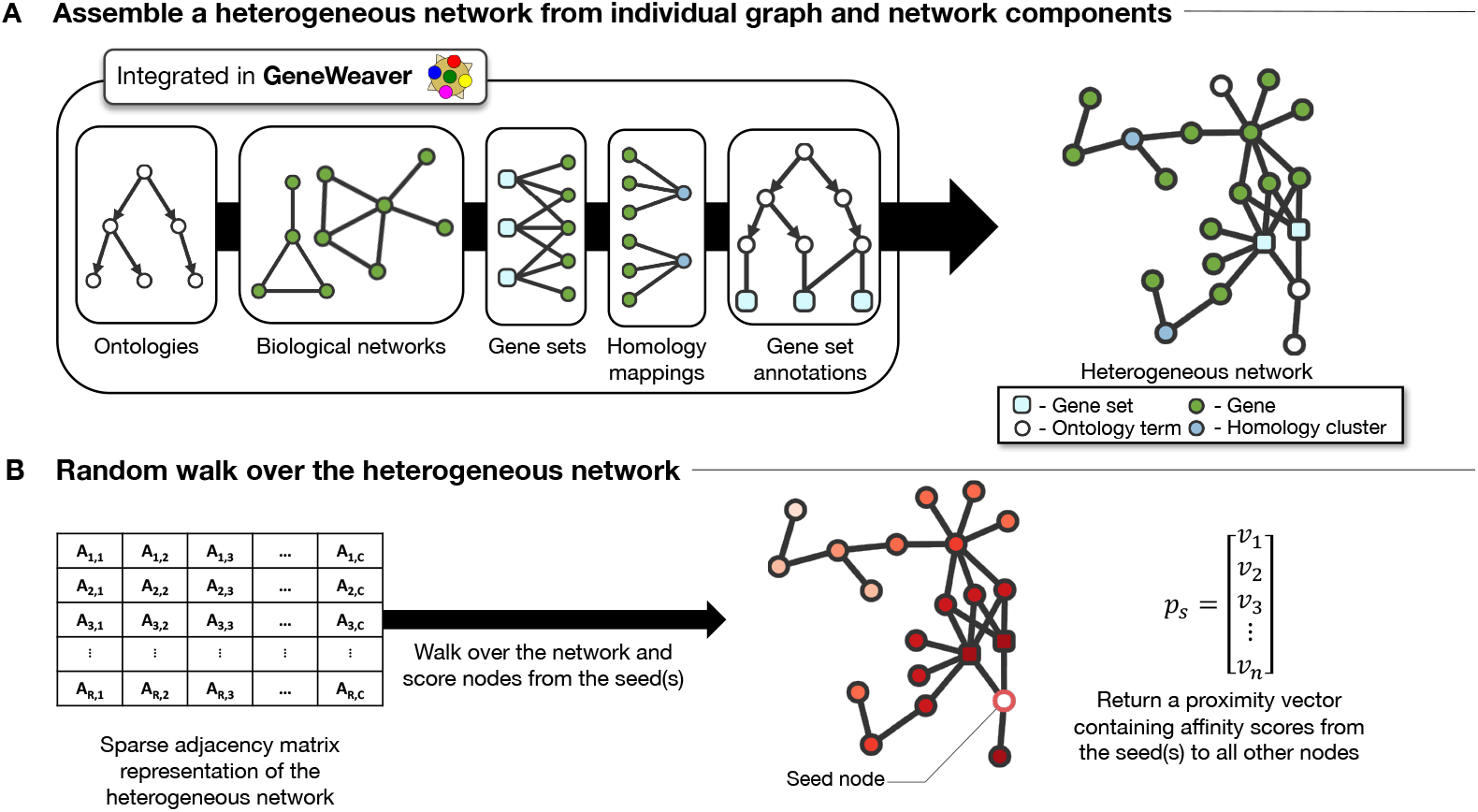
NESS heterogeneous network construction and graph walk. A multi-species network comprised of functional genomics and semantic data is built from several data graphs including ontological relationships and annotations, biological networks, set-gene associations, and homology clusters (**a**). After network construction, similarities between entities in the graph are enumerated using a random walk with restart (**b**).

**Table 1.**
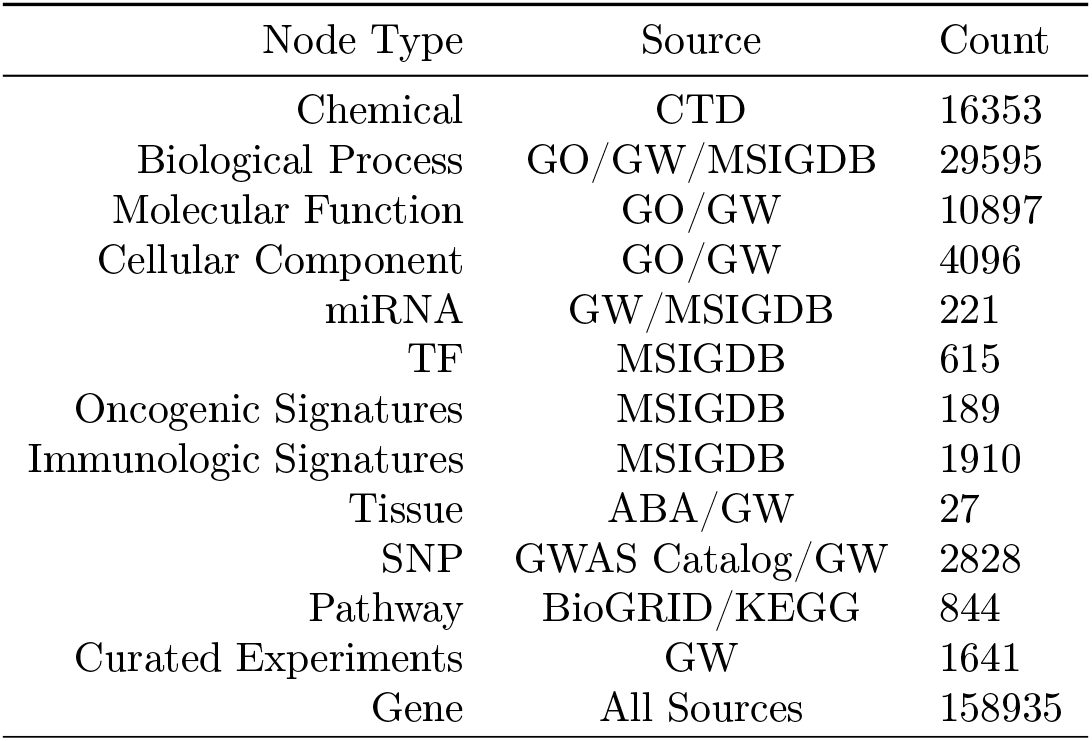
Node types within the heterogeneous network, their data source, and the number of nodes for each type present in the final network. Abbreviations: Allen Brain Atlas (ABA), Comparative Toxicogenomics Database (CTD), Gene Ontology (GO), GeneWeaver (GW), Kyoto Encyclopedia of Genes and Genomes (KEGG), Molecular Signatures Database (MSIGDB).

Many of the data sources used by NESS are also available in GeneWeaver for derived analyses. GeneWeaver (https://geneweaver.org) is a suite of services that function as a multiple-species, heterogeneous data store and analysis platform [25]. It is designed to integrate a variety of experimentally driven genomic data types including differential expression profiling, genome-wide association studies (GWAS), literature curated data, and gene coexpression results for ten different species. In addition to curated experimental datasets in GeneWeaver, data available for analysis includes extensive public resources such as pathway and network databases (e.g., Pathway Commons), ontology annotations from the Mouse Genome Database (MGD), Rat Genome Database (RGD), etc. (e.g., GO, HP). It also includes various curated sources (e.g., Comparative Toxicogenomics Database, GWAS Catalog, Allen Brain Atlas). The inclusion of public resource data, particularly those defining biological states, aids in the discovery of genes shared between biological processes and disease phenotypes across species. NESS uses GeneWeaver’s bipartite data model as one of many graph-based integrations.

Using the collected networks, curated gene sets, and public resource data, a large graph of biological entities was constructed. Directed, heterogeneous networks were built for *Homo sapiens*, *Mus musculus*, and *Rattus norvegicus* using species specific data sets. An additional network incorporating homologous genes conserved across the three species was also generated.

### Preparation of integrated data resources

#### Gene Ontology annotations

GO structures and annotations were retrieved (November 2018) for *H. sapiens, M. musculus*, and *R. norvegicus*. Relationships that were not “is a”, “part of”, or “regulates” (including positvely and negatively subtypes) were removed from the GO graph. All electronically inferred annotations (IEA) were excluded. Only experimental (EXP, IDA, IEP, IGI, IMP, IPI), author statement (TAS), or curatorial statement (IC) evidence is used. The exclusion of IEA evidence is done to remove inferred orthology-based annotations and ensure only the highest quality annotations are used—primarily those reviewed and validated by expert curators.

#### Interaction networks

Interaction data from two biological network resources, the Kyoto Encyclopedia of Genes and Genomes (KEGG) [26] and BioGRID [27], were retrieved (November 2018). In total, 746 pathways from KEGG is used, spanning the following categories: “Metabolism”, “Genetic Information Processing”, “Environment Information Processing”, “Cellular Processes”, and “Organismal Systems”. Curated KEGG pathways are used as a ground truth for metric comparison since pathways in these categories contain gene networks for each species used in this study. Two categories, “Human Disease” and “Drug Development” were excluded since these categories contain pathways specific to a single organism or do not have any usable gene-gene associations. Protein-protein interactions (PPI) from BioGRID for each of the three species were downloaded. PPI networks were reconstructed and incorporated with the remaining network data sets.

#### GeneWeaver data sets

Curated, experimental gene sets were retrieved from GeneWeaver (November 2018). Each of these sets met the following criteria: i) a size of less than 5000 genes and greater than 1 gene, ii) derived from published, publicly available research, and iii) were not positional candidate gene sets or the results of quantitative trait loci (QTL) analysis and mapping. This provided an additional 1641 gene sets with an average of 88 genes per set. In addition, GeneWeaver typically integrates several public resources which were also included in this study: 16,353 curated chemical-gene interaction data sets from the Comparative Toxicogenomics Database (CTD) [28], 27 gene sets from the Allen Brain Atlas (ABA) [29] examining differential expression patterns in the adult mouse brain, and 2844 human SNP-trait association gene sets collected from the GWAS Catalog [30]. Also included were 221 sets of miRNA targets, 615 sets whose genes contain regulatory motifs that function as transcription factor (TF) binding sites, 189 sets containing pathway genes disrupted in cancer, 1910 sets containing genes associated with immune system processes, and 50 sets representing well defined biological processes, all of which are integrated from the Molecular Signatures Database v.6.0 (MSigDB) [31].

#### Species integration

The data was partitioned by species and an additional group of sets incorporating homologous genes between species was constructed using GeneWeaver’s store of homology associations. GeneWeaver uses its own dynamic internal identifiers, known as cluster IDs, to map multiple gene identifiers to a single canonical gene entity. Canonical genes are defined by each model organism database (MOD) and used as the primary gene reference for that species. In the absence of an established MOD, NCBI Entrez is used. This allows GeneWeaver to unify protein isoforms, gene transcripts, genetic variants, and alternate gene references or synonyms into a single unique identifier: a GeneWeaver ID (GWID). After the generation of canonical gene groups, cross-species associations are produced by mapping GWIDs (and by extension, gene groups) to homology clusters retrieved from NCBI Homologene [32] and MGI [33]. During the data collection and integration process, any genes that could not be confidently mapped to a single GeneWeaver ID or ortholog cluster were excluded.

### NESS: graph walks over heterogeneous networks

To determine similarity among terms and other entities, a random walk with restart (RWR) is used [34]. The RWR iteratively traverses the graph from a start node or nodes referred to as seed(s) and results in a probability distribution representing the likelihood of visiting a particular node when originating from the seed(s). This metric can be thought of as the affinity between nodes in the graph. At each iteration of the walk, the algorithm will either a) randomly visit a new node selected from the neighbors of the current node or b) restart (using a predefined restart probability) from the seed(s). Higher restart probabilities force the walk to assess the local neighborhood which surrounds the seed(s), while lower restart probabilities cause the algorithm to examine the global topology of the graph.

*G* = (*V, E*) is a directed graph with a set of vertices, *V* = {1, 2, …, *n*} and a set of edges, *E* ⊂ *V* × *V*. Given a directed graph *G* in the form of a column normalized adjacency matrix *A*, the random walk among the graph and proximity values for a specific node *s* can be defined as

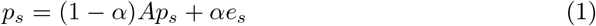

where *α* is the restart probability; *p_s_* is the proximity vector of node *u*; *e_s_* is the initial proximity vector in which *e_s_*(*s*) = 1 and all other values are 0 in the case of a single seed node, or equal probabilities in the case of multiple start nodes.

This formulation of the walk allows *p_s_* to be iteratively calculated using matrix and vector multiplications until a steady state is reached. The algorithm converges once the *L*_1_ norm of the difference between proximity vectors at two successive iterations reaches a sufficiently low value:

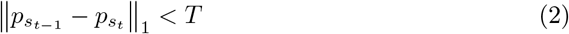

where *t* is the iteration and *T* is a small number such as 10^−8^ which serves as a convergence threshold.

The final result of the walk is a proximity vector containing affinities from some node *s* to all other nodes in the graph. Unless otherwise specified, a restart probability of 0.35 and a convergence threshold of 10^−8^ was used for every random walk in this analysis.

A heterogeneous network, represented as an adjacency matrix, is built iteratively from the various graphical data representations retrieved from GeneWeaver (Algorithm 1). The random walk is used to assess the affinity between two entities, ontology terms for example, in the graph (Algorithm 2). NESS, the combined network construction and random walk, enable semantic comparisons infused with a rich set of functional genomics data (Fig 1).

**Algorithm 1:**
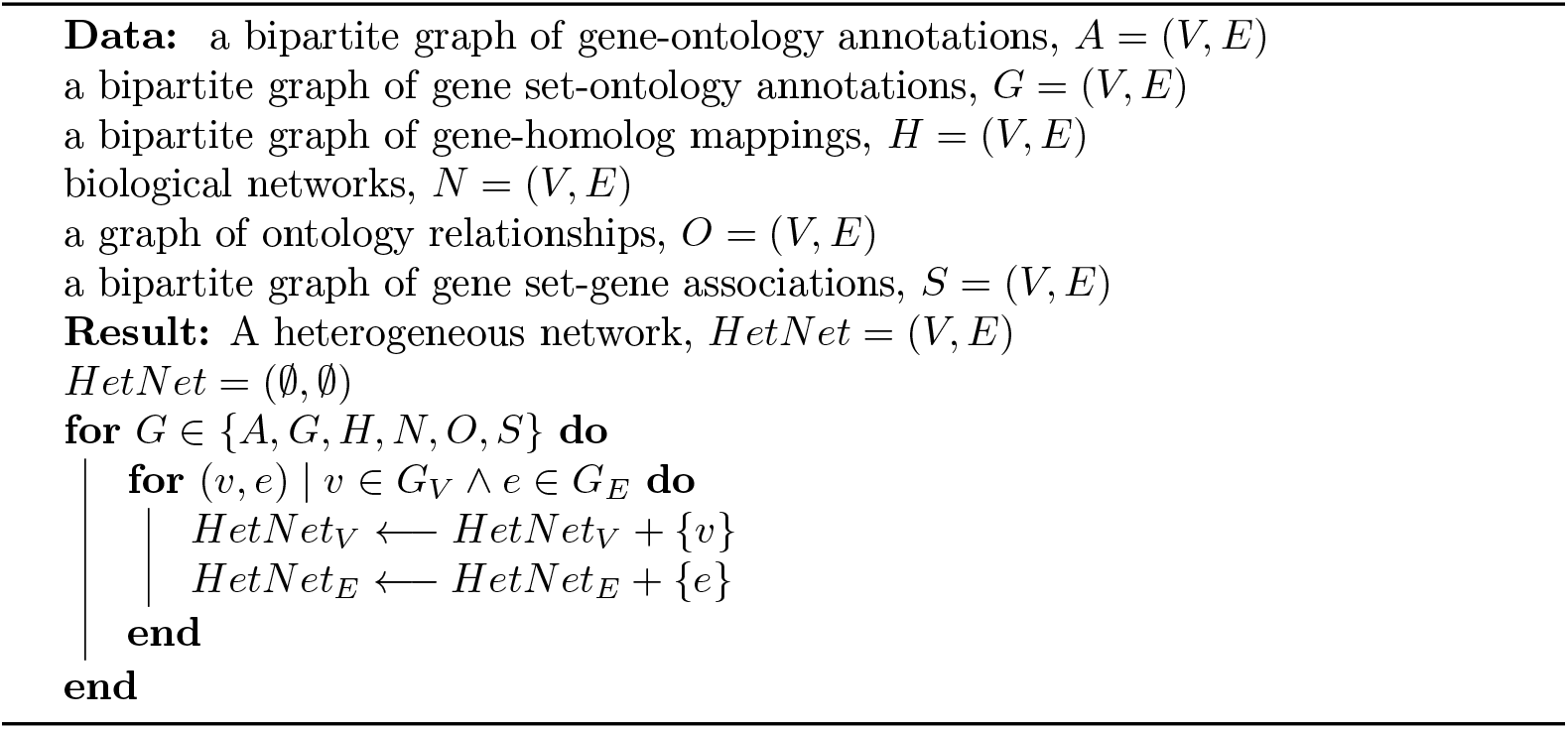
Heterogeneous network assembly

**Algorithm 2:**
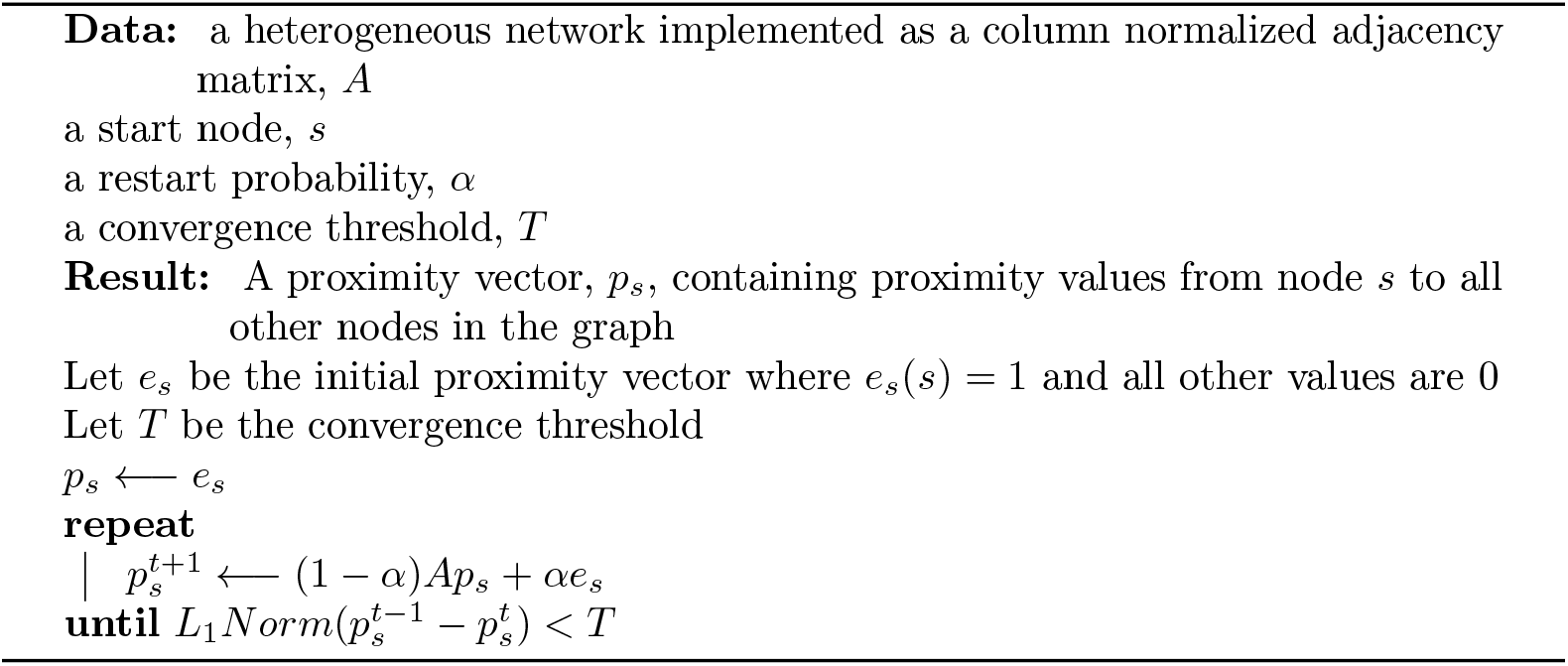
Random walk with restart over a heterogeneous network

### Similarity metric evaluation

To evaluate the performance of NESS in relation to other similarity methods for term comparison, objective criteria is needed. However, as no gold standard criteria exists, the best available option is to use prior knowledge as a ground truth for similarity metric comparison. Curated functional genomic pathways provide a robust context for this evaluation. Gene products are well characterized and annotated. These annotations can be leveraged to compare the functional similarity of genes in order to develop an unbiased metric with which to compare semantic and data-centric similarity methodologies.

To determine the similarity among gene products, we adapt the functional similarity methodology from [35]. The similarity between two genes is computed using maximum semantic similarity scores from the annotation sets of both genes:

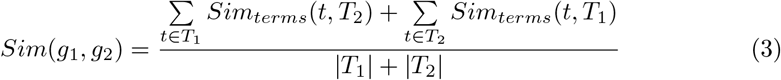

where *g*_1_ and *g*_2_ are two genes; *T*_1_ and *T*_2_ are the set of terms annotated to genes *g*_1_ and *g*_2_, respectively. The similarity between a term and a set of terms, *Sim_terms_*, is defined as:

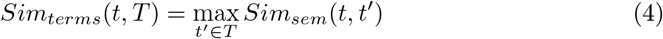

where *Sim_sem_* can be any known similarity measure. For our purposes, *Sim_sem_* is replaced with Resnik, Lin, Jaccard, and cosine measures.

Our criterion for assessing and comparing the accuracy of similarity metrics is based on prior knowledge in the form of curated biological pathways. The biological function of processes shared by genes within the same pathway exhibit greater cohesiveness than that of genes in separate pathways. Therefore, gene similarities can be examined as a ratio of intra-pathway and inter-pathway relatedness, resulting in a pathway score, *PS*:

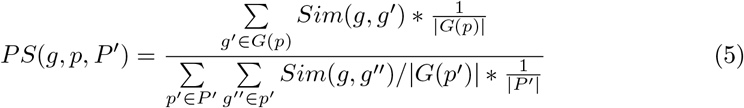

where *p* is some pathway, *P*’ is a set of pathways which does not contain *p*, and *G*(*p*) is the set of genes in pathway *p*. To prevent bias, comparisons between two pathways which share genes are not considered.

### Prioritization of gene-disease associations and classification of disease-associated gene function

To assess the utility of NESS in identifying and prioritizing genes associated with complex disease, prioritized genes were compared to “gold standard” gene sets retrieved (June 2019) from DisGeNET [36], a resource for curated and mined gene-disease associations. Gene prioritization assessments were conducted for a subset of substance use disorders: alcohol, heroin, morphine, and nicotine dependence. For each gene prioritization test, the ground truth gene set from DisGeNET was divided evenly into seed and test sets. Genes in the seed set were used to seed NESS, and its ability to recapitulate genes in the test set was measured. Gene prioritization was also assessed and compared with several other methods using this same approach. Other state-of-the-art methods tested include a random walk using only interaction datasets [14], a combinatorial approach based on the enumeration of maximal biclique modules (HiSim) [25], and a disease module detection approach using the human interactome (DIAMOnD) [12].

#### Testing gene-disease associations produced by random walks

Gene-disease associations were prioritized using random walks over interaction datasets, in a methodolgy similar to the one specified in [14]. Human interaction datasets from KEGG and BioGRID were used as input. A restart parameter, *α* = 0.35, was used for the random walk. Prioritized genes with a random walk score of at least 5%, *RWR_score_* >= 0.05, and p-value, *p* < 0.01, after graph permutation testing were compared to ground truth datasets.

#### Testing gene-disease associations produced by maximal biclique modules

Genes were prioritized using GeneWeaver’s Hierarchical Similarity (HiSim) graph tool [25]. The HiSim graph tool enumerates maximal bicliques from gene set-gene bipartite graphs and constructs a directed acyclic graph (DAG) from overlapping bicliques. Maximal modules within the DAG (*i.e.,* nodes lacking any parents) contain prioritized genes shared among many gene sets. Although not a strict one-to-one comparison with other methods, use of the HiSim graph tool tests state-of-the-art combinatorial approaches to gene prioritization.

Input for the HiSim graph tool are collections of public gene sets derived from published, functional genomics experiments. Two separate HiSim tests were performed in GeneWeaver: one utilizing only human datasets, and another which makes use of GeneWeaver’s cross-species analysis capabilities to include *M. musculus* and *R. norvegicus* gene sets assessing chronic and acute drug use. Genes present in maximal HiSim modules were compared to ground truth datasets from DisGeNET.

#### Testing gene-disease associations produced by disease module detection

A DIseAse MOdule Detection (DIAMOnD) algorithm [12] iteratively builds gene modules associated with disease using gene interaction networks. Given a set of seed genes, DIAMOnD grows the module by identifying significant connectivity patterns at each iteration. Testing used default parameters except for the output module size which was set to 250–slightly larger than the default setting to ensure all genes from the testing sets could be recapitulated. Network inputs included BioGRID, KEGG, and STRING. STRING-based interactions were added to replicate the curated, physical, high-throughput, and literature-mined protein-protein associations used in the original DIAMOnD study. Genes in DIAMOnD-produced modules were compared to ground truth DisGeNET sets.

### Implementation

A fast, scalable implementation of NESS is provided, written in Python 3.7, which makes use of optimized numerical analysis libraries for better performance. Additionally, this implementation is also designed for use with big data; NESS has built-in support for high performance computing and cloud environments, and we demonstrate that its performance scales with increased CPU cores (Fig S1). Finally, NESS has been integrated into the GeneWeaver resource, which allows realtime analysis of functional genomics datasets via a web interface.

## Results

### Network Enhanced Similarity Search performance evaluation

NESS performance was evaluated through two distinct characterizations: first, as a metric for comparing biological entities modeled using ontologies, and second, via its utility for prioritizing genes associated with complex disease. In both instances, expert curated resources function as a ground truth.

#### Applying NESS to comparisons of biological entities

To evaluate the accuracy and precision of term similarity estimated by the NESS heterogeneous graph and random walk model, we examined the performance of NESS in comparison to well-established similarity measures and using KEGG as a ground truth data set. Each term similarity metric was evaluated under several conditions. The *H. sapiens*, *M. musculus*, and *R. norvegicus* datasets were evaluated separately to measure the impact of the NESS RWR aspect of the analysis without the complication of multiple species. An independent test using a full implementation of NESS, which includes an aggregated multi-species graph, was used to evaluate the impact of heterogeneous data sets on similarity accuracy and precision.

Within individual species, and under conditions optimized for the Resnik, Lin, Jaccard, and cosine similarity metrics, the walk-based NESS approach produced significantly better results across species specific datasets than Resnik, Lin, Jaccard, or cosine measures (Fig 2), with an AUC of 0.89, 0.74, and 0.68 for *H. sapiens, M. musculus*, and *R. norvegicus*, respectively (Fig 3). Comparisons of the AUCs using Mann-Whitney U indicates these results include a 25 percent increase over Resnik (*p* < 0.001), a 30 percent increase over Lin (*p* < 0.001), a 19 percent increase over Jaccard (*p* < 0.001), and a 19 percent increase over Cosine (*p* < 0.001) for *H. sapiens*; a 38 percent increase over Resnik (*p* < 0.001), a 47 increase over Lin (*p* < 0.001), a 38 percent increase over Jaccard (*p* < 0.001), and a 38 percent increase over cosine (*p* < 0.001) metrics for *M. musculus*; and a 35 percent increase over Resnik (*p* < 0.001), a 36 percent increase over Lin (*p* < 0.001), a 36 percent increase over Jaccard (*p* < 0.001), and a 36 percent increase over cosine (*p* < 0.001) for *R. norvegicus* specific data. Resnik produced slightly better results than Lin for each of the three species–a result that is consistent with previous similarity comparisons [19].

**Figure 2.**
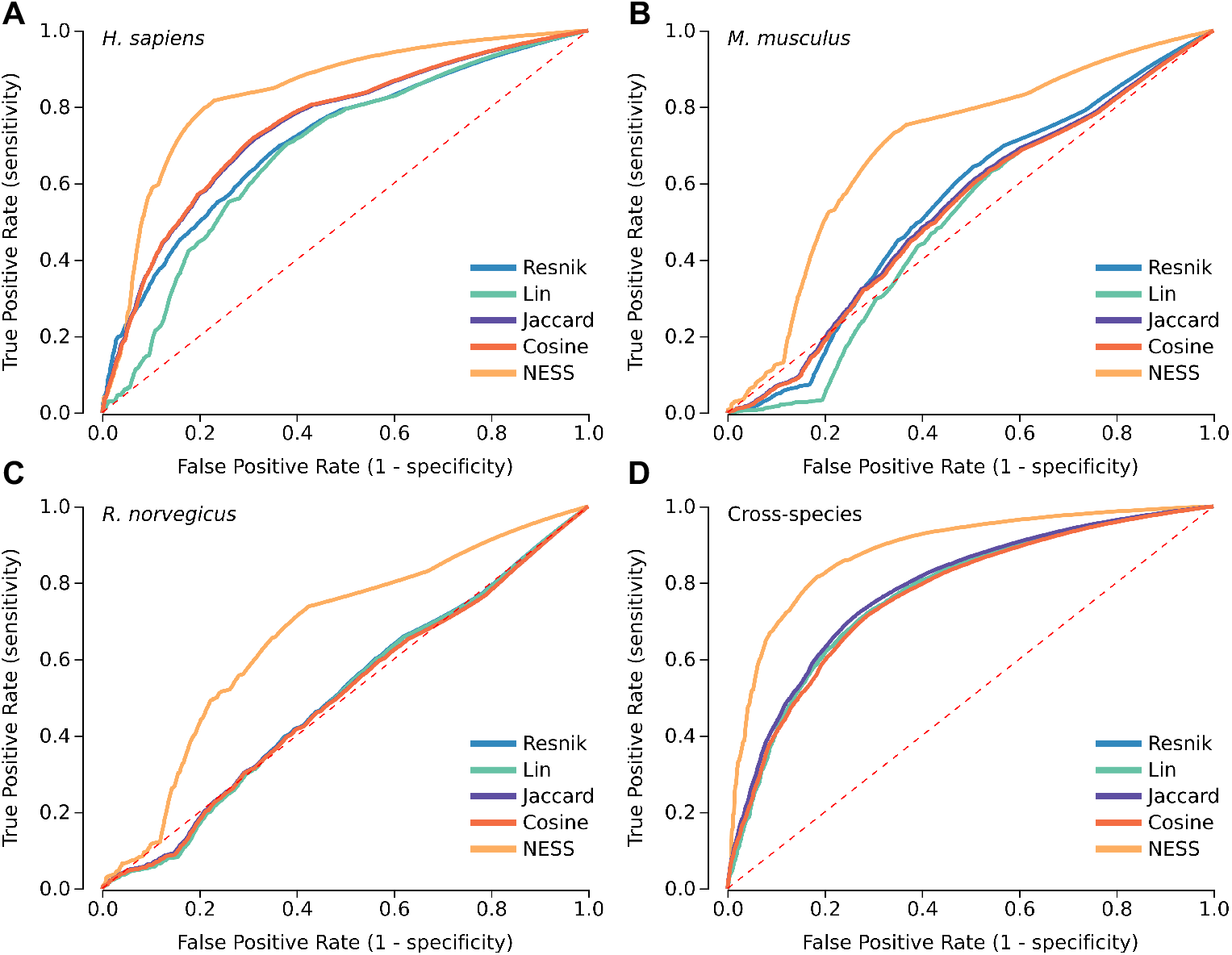
ROC curves comparing similarity metric performance of Resnik, Lin, Jaccard, cosine, and NESS based measures. Gene function in curated KEGG pathways for *H. sapiens* (**a**), *M. musculus* (**b**), *R. norvegicus* (**c**), and cross-species (**d**) data sets were used as a ground truth.

**Figure 3.**
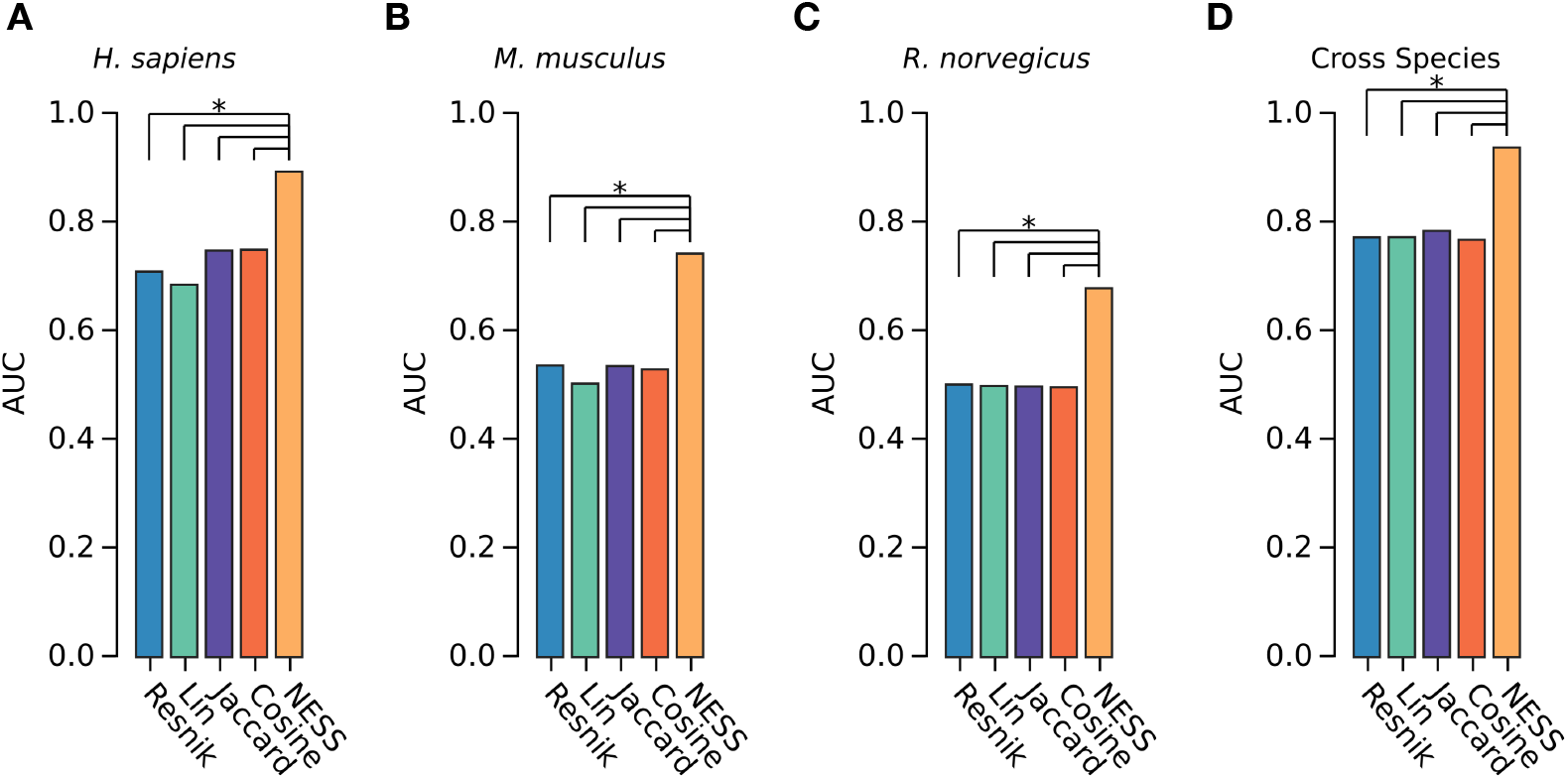
AUC scores for each similarity metric and species in the study. Plots show significant differences in the AUC produced by NESS compared to Resnik, Lin, Jaccard, and cosine metrics. Significance of differences among AUCs was determined using Mann-Whitney U (∗*p* < 0.001).

Discovery rates of the ground truth KEGG data demonstrates that the traditional semantic similarity metrics of Resnik and Lin performed better when the data includes homologous gene associations relative to a single species. The NESS approach demonstrated a significant improvement over all single species data sets with an AUC of compared to an AUC of 0.64 (*p* < 0.001) for Resnik, an AUC of 0.71 (*p* < 0.001) for Lin, an AUC of 0.78 for Jaccard (*p* < 0.001), and an AUC of 0.76 for cosine (*p* < 0.001) similarity (Fig 3d). Importantly, all methods that use aggregated cross-species graph data perform better than single species data sets alone. F1 scores (Fig S2) and a complete confusion matrix (Table S1) for each of these tests reiterate that the performance of NESS generates greater accuracy, precision, recall and specificity across each comparison.

#### Applying NESS to the classification of gene function in genomic studies of disease

NESS was applied to a collection of cross-species functional genomics data (Table S2) to assess its utility in recapitulating complex disease-related gene sets. This was approached by evaluating the similarity of curated gene disease associations with those derived from aggregated genome wide experimentation. Certain groups of complex disease, such as substance use disorders (SUD), are difficult to classify and study due to their heterogeneity, comorbidity, and symptom overlap with other conditions [20]. NESS was used to refine and classify gene-disease associations by leveraging animal experiments which assess dependence and consumption phenotypes, biological networks, and curated data stores. Performance was compared to other approaches including data-specific simple random walks [14], combinatorial prioritization using GeneWeaver’s Hierarchical Similarity (HiSim) graph tool [25], and DIsease MOdule Detection (DIAMOnD) [12]. On average, NESS exhibited increased performance at recapitulating known disease-gene associations for SUDs than other prioritization approaches (Fig 4a – d). Instances where few disease-specific datasets exist, and biological knowledge is primarily relegated to gene interaction networks, DIAMOnD had comparable performance to NESS (Fig 4b,c). Among the tested methods, HiSim was the only combinatorial approach to prioritization. This approach relies on an abundance of input datasets related to specific conditions, phenotypes, or diseases. These results indicate that in cases where human data is sparse, animal model studies can aid the prioritization of genes associated with human disease; in all test cases, the cross-species HiSim (CS HiSim) results recapitulated more disease-gene associations than human datasets alone. Additionally, NESS also demonstrated improved performance when comparing SUD experimental studies to their respective “gold standard” gene sets (Fig 4e), and by assessing the proportion of significant similarity scores among SUD experiments and curated annotations (Fig S3).

**Figure 4.**
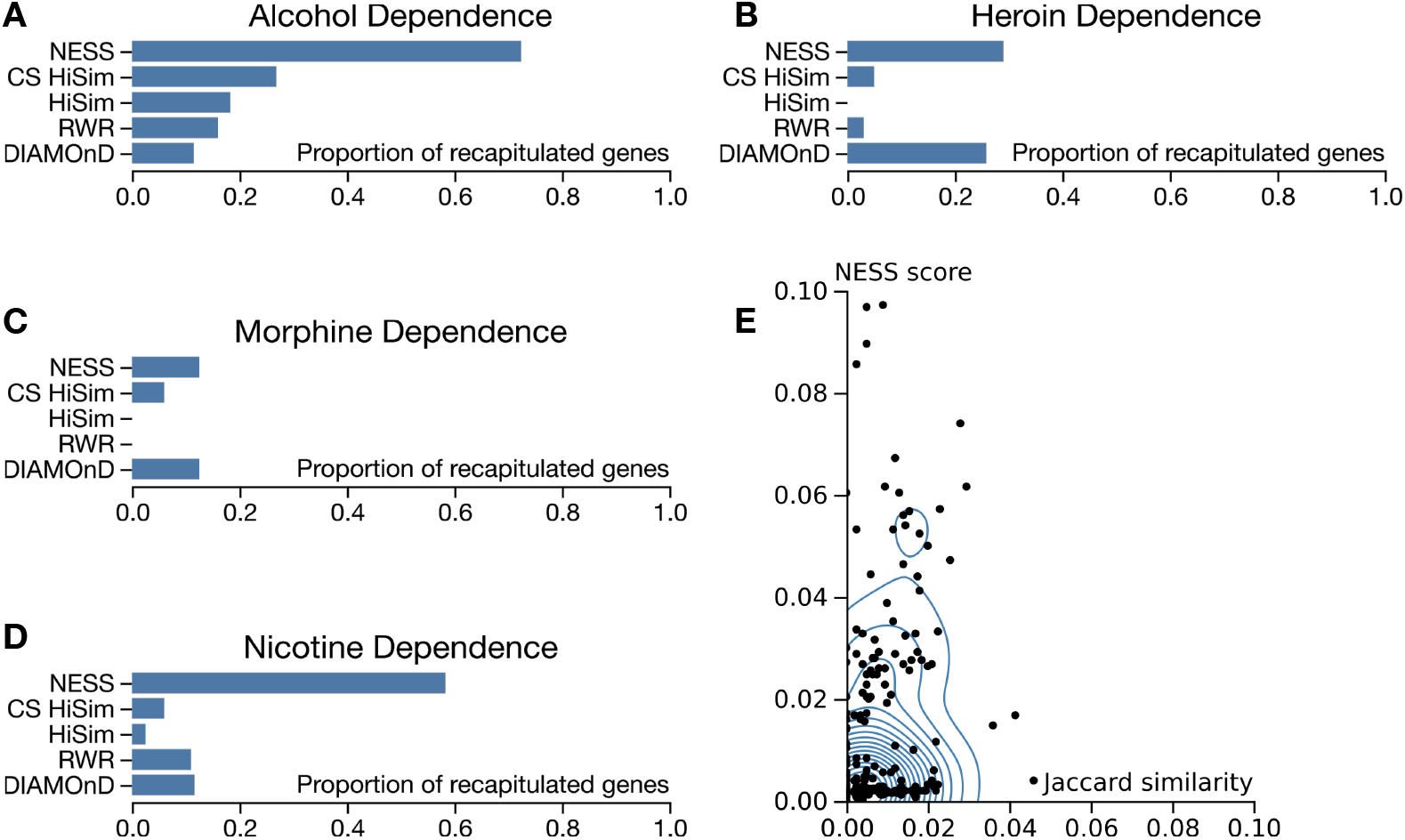
Recapitulating known disease-gene associations from functional genomics data. A single heterogeneous graph was built from substance use disorder (SUD) experimental studies from GeneWeaver, the Disease Ontology (DO), BioGRID protein interactions, and KEGG pathways. Known disease-gene associations from DisGeNET were witheld from the graph integration and used a “gold standard” dataset for comparison. Disease-gene associations produced by NESS were only used if their normalized NESS probability was at least 5% (*p_NESS_* >= 0.05) and the association was significant after permutation testing (*p* < 0.01, *n* = 2500). NESS recapitulated a higher amount of curated substance use disorder gene annotations than set overlap techniques (**a** - **d**). Additionally, NESS scores among SUD experimental studies and their respective “gold standard” datasets were higher than Jaccard coefficients for the same comparison (**e**). Abbreviations: DIseAse MOdule Detection (DIAMOnD), Hierarchical Similarity (HiSim), Cross-species Hierarchical Similarity (CS HiSim), random walk with restart (RWR), Network Enhanced Similarity Search (NESS).

#### Applying NESS to identify genes involved in multiple substance-use disorders

To demonstrate one application of NESS, novel disease-gene associations were identified and classified as multiple or single SUD from functional genomics data. Using a two-fold threshold of a NESS probability score of 1% (*p_NESS_* >= 0.01) and a p-value threshold of *p* < 0.01, 400 novel, putative SUD-gene associations were identified (Table S3). Seven of these genes were associated with at least three SUDs, indicative of their potential association with polysubstance abuse. Gene Ontology enrichment analysis (Table S4) revealed these genes to be enriched in catecholamine metabolic processes (GO:0006584, *p* = 2.4 · 10^−6^), arachidonic acid secretion (GO:0050482, *p* = 7.7 · 10^−5^) and transport (GO:1903963, *p* = 7.7 · 10^−5^). Serotonin receptor (*HTR2A*) and transport (*SLC6A4*) genes were also present among multiple SUD associations for alcohol, heroin, and nicotine dependence. These genes are well studied for their roles in the development of single substance use disorders [21] but only recently has their role in simultaneous alcohol and heroin dependence been investigated [22]. Perturbations in archidonic acid processes also represent an interesting avenue of study in drug addiction due to their role in often co-occurring psychiatric disorders, namely bipolar disorder [23]. Overall, these results illustrate the utility of NESS in gene function classification and elucidating the genomic basis of complex disease.

### Resilience of Network Enhanced Similarity Search to sparse and noisy data

A primary motivation for using a walk-based approach on heterogeneous networks to augment similarity measurements is to prevent biases in results due to noisy or missing data. This approach was tested by simulating missing data by removing random edges from the graph. Using the same input and evaluation criteria as specified in the methods for comparing ontology terms, 0.5%, 1%, 2.5%, 5%, 10%, 15%, 20%, 25%, and 30% of the graph’s total edges were removed and AUC was reevaluated. Results illustrate that missing relationships did not substantially impact AUC scores up to 30% (1,358,520) of the graph’s total edges (Fig 5a). Likewise, noisy data sets simulated by adding percentages of random edges had similar effects on the final AUC scores (Fig 5b). These results indicate that the NESS method works well in the case of sparse or noisy datasets.

**Figure 5.**
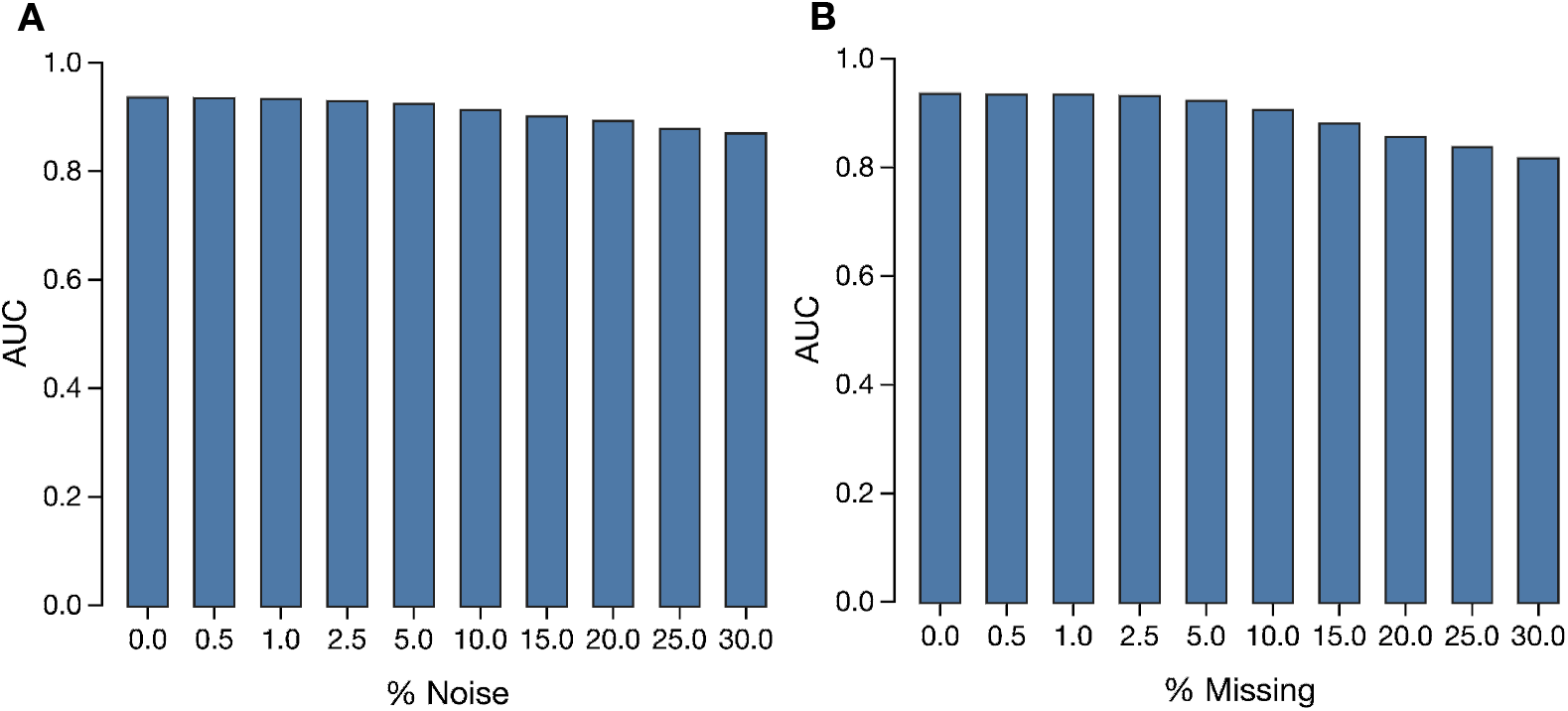
Evaluating graph walk robustness. Graphs are built using all available resources, the Gene Ontology’s (GO) Biological Process (BP) ontology, and homology data sets. The restart probability, *α* = 0.35 was used. Missing data was simulated by randomly removing 0.5%, 1%, 2.5%, 5%, 10%, 15%, 20%, 25%, and 30% of the graph’s total edges which accounts for 22642, 45284, 113210, 226420, 452840, 905680, 1132100 679260, and 1358520 edges respectively. Noisy data was simulated by randomly adding new edges in 0.5%, 1%, 2.5%, 5%, 10%, 15%, 20%, 25%, and 30% proportions of the graph’s total edges. Despite missing or false associations, the performance of the graph walk remains constant as indicated by area under the curve (AUC) measurements.

### Optimization of Random Walk Restart Probability

We evaluated the NESS algorithm to determine the optimal restart probability for the heterogeneous data sets used in this study. Higher restart probabilities force the walk to analyze the local topology of the graph rather than its global layout. The same input and evaluation criteria for comparing ontology terms was used to test restart parameters. After examining eight different restart probabilities, *r* = 0.35 was determined to be optimal and this parameter was used for each analysis in this study (Fig 6).

**Figure 6.**
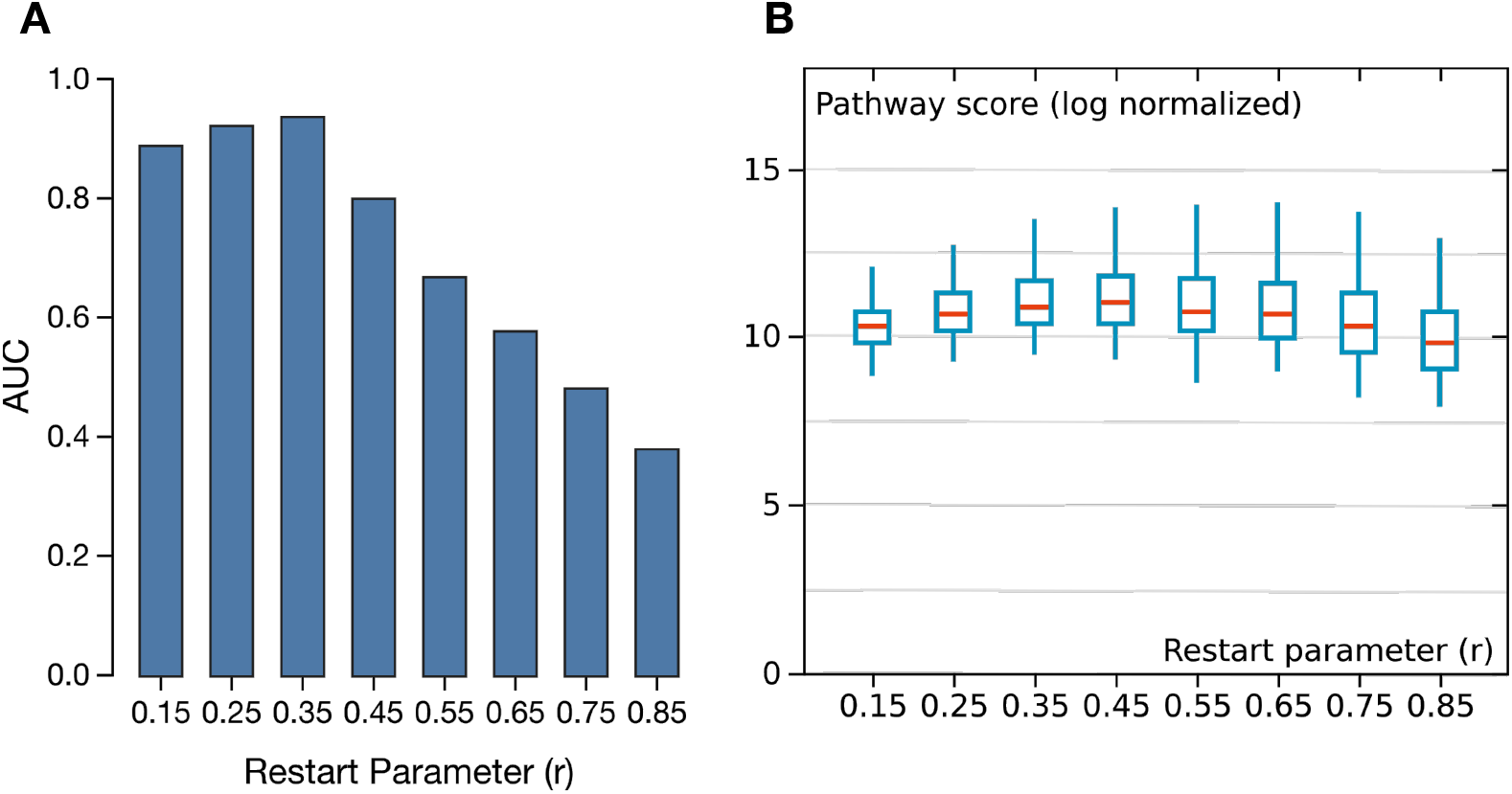
Evaluating optimal RWR restart parameters. Graphs are built using all available resources, the Gene Ontology’s (GO) Biological Process (BP) ontology, and homology data sets. Eight different restart probabilities are tested. AUC and pathway scores indicate that *α* = 0.35 is the optimal restart probability given the data types used as input (**a**, **b**).

## Discussion

In the present study we designed and tested a methodology to leverage large-scale functional genomics data from GeneWeaver to improve concept-based similarity analysis. We show this method is effective for comparing ontology terms, even among sparsely annotated concepts, and we developed and examined the effectiveness of NESS, a methodology for improving similarity metrics through the inclusion and analysis of cross-species, functional genomics data. Results against known data illustrate that NESS improves sensitivity and accuracy for assessing the similarity between ontology concepts. For the GO, NESS demonstrated significantly better performance than similarity measures. These measures were selected based on their performance in previous comparative studies [19]. Our results illustrate that the harmonization of functional genomics data and homologous relationships, coupled with the analysis of graph properties, results in an improved similarity metric. Additionally, NESS is effective at handling sparse or spurious datasets–useful for studying concepts lacking annotations or those with noisy associations. These findings indicate that the aggregation and use of data from multiple experimental assays, high-throughput studies, and evolutionary relationships can be used to significantly improve concept comparison and alleviate species or assay-specific biases from annotations.

Although NESS can be used as an enhanced similarity measure for biological concepts, it is not restricted to concept-concept comparisons. The flexible graph model and RWR algorithm allow any pair of biological entities to be associated, such as genes, variants, or regulatory mechanisms. Node and edge types can be automatically inferred based on metadata and annotations associated with the integrated data source. Gene-based entities can be further refined using GeneWeaver’s internal representation of gene types which includes support for SNPs, proteins, non-coding transcripts, and regulatory elements. Any biological component that can be associated with or rolled up to the gene level can be included in the graph. For example, variants that may reside in or regulate a gene can be associated with that gene for analytical purposes. Such a tool has applications in the interpretation of genomic data in light of model organism data and the prioritization of genes and variants. Gene or variant prioritization can assess the contribution of genetic effects to a particular disease process. Indeed, random walks have previously seen success in both disease-gene discovery [14] and variant prioritization [24], but have not been used for these purposes against the background of complex datasets utilized by NESS. More recently, complex empirical datasets have been used in conjuction with network mining techniques to prioritize gene-disease associations [12, 16, 17]. While effective, these recent improvements have only utilized a single data type (e.g., gene networks) or species, despite an array of functional genomics and animal model studies available for use. Our work confirms that additional empirical evidence is useful for refining gene-disease associations and shows that data aggregation from multiple disparate resources, systems, and model organism studies can be a powerful component of gene-disease prioritization.

Finally, examining disease similarity using genomic data as a basis for comparison can help elucidate the molecular mechanisms involved in seemingly disparate disease processes. We illustrate examples in which NESS exhibited improved performance in prioritizing disease-gene associations compared to other prioritization methods. We also identified novel gene-SUD associations enriched in archidonic acid secretion and transport. Changes in these biological processes have been well documented in other co-occuring disorders [23]. Additionally, genes involved in serotonergic pathways were identified across multiple dependence disorders. Although these genes are well studied in alcohol dependence, these findings suggest they have a greater role in other SUDs. Overall, these applications make NESS a useful tool for genome-level analysis. We make this tool freely available for use within the GeneWeaver resource https://geneweaver.org, and its source code available for download and use at https://github.com/treynr/ness.

## Acknowledgments

The authors thank Stephen Krasinski for his helpful review and comments.

## Supporting information

**Figure S1 Scalability of NESS for big datasets.** NESS performance and scalability was tested using a randomly generated, scale-free graph consisting of 10,000 nodes with a density, *D* = 0.1. The amount of time taken to calculate the proximity matrix, *P_M_* (100,000,000 comparisons), was determined using the average of four separate NESS runs. Average runtime, measured in seconds, decreases as additional cores are used by NESS. All tests were conducted on identical systems containing Intel Xeon E5-2695 CPUs @ 2.10GHz.

**Figure S2 F1 scores evaluating the performance of Resnik, Lin, Jaccard, cosine, and NESS measures for *H. sapiens*, *M. musculus*, *R. norvegicus*.** Curated KEGG pathways are used as a ground truth for similarity metric comparison.

**Figure S3 Significant similarity scores among substance use disorder (SUD) experimental studies and “gold standard” datasets comprised of curated disease-gene associations.** SUD experimental studies were compared to gold standard datasets using NESS and Jaccard similarity. Curated disease-gene associations were retrieved from DisGeNET. Significant Jaccard coefficients were determined by generating a null distribution of coefficients using premutation testing (n = 2500). NESS score significance was assessed by permuting graph labels and recalculating the walk statistic (n = 2500). Overall, the proportion of significantly similar experimental SUD studies to their respective gold standard dataset was higher when using NESS.

**Table S1 Confusion matrices assessing the performance of Resnik, Lin, Jaccard, cosine, and NESS measures at recapitulating ground truth data from KEGG.**

**Table S2 Experimental gene sets from substance dependence studies curated in GeneWeaver and used to assess the performance of NESS at classifying complex disease.**

**Table S3 Novel substance dependence associated genes prioritized using NESS.**

**Table S4 GO biological process enrichment analysis of substance dependence genes prioritized using NESS.**

## Author Contributions

TR, EJC, and EJB conceived and designed the experiments. JAB curated experimental datasets. TR, EJB, and MAL implemented data models and analysis tools. TR analyzed the data. TR, EJB, and EJC wrote the paper. EJB and EJC carried out manuscript revisions.

